# Pulmonary Metagenomic Sequencing Suggests Missed Infections in Immunocompromised Children

**DOI:** 10.1101/291864

**Authors:** MS Zinter, CC Dvorak, MY Mayday, K Iwanaga, NP Ly, ME McGarry, GD Church, LE Faricy, CM Rowan, JR Hume, ME Steiner, ED Crawford, C Langelier, K Kalantar, ED Chow, S Miller, K Shimano, A Melton, GA Yanik, A Sapru, JL DeRisi

**Author notes:** **CONTACT**: Matt Zinter, MD, University of California - San Francisco, Department of Pediatrics, Division of Critical Care Medicine, 550 16^th^ St, San Francisco, CA 94143, USA.

## Abstract

**RATIONALE:** Despite improved diagnostics, pulmonary pathogens in immunocompromised children frequently evade detection, leading to significant morbidity and mortality.

**OBJECTIVES:** To develop a highly sensitive metagenomic next generation sequencing (mNGS) assay capable of evaluating the pulmonary microbiome and identifying diverse pathogens in the lungs of immunocompromised children.

**METHODS:** We collected 41 lower respiratory specimens from 34 immunocompromised children undergoing evaluation for pulmonary disease at 3 children’s hospitals from 2014-2016. Samples underwent mechanical homogenization, paired RNA/DNA extraction, and metagenomic sequencing. Sequencing reads were aligned to the NCBI nucleotide reference database to determine taxonomic identities. Statistical outliers were determined based on abundance within each sample and relative to other samples in the cohort.

**MEASUREMENTS & MAIN RESULTS:** We identified a rich cross-domain pulmonary microbiome containing bacteria, fungi, RNA viruses, and DNA viruses in each patient. Potentially pathogenic bacteria were ubiquitous among samples but could be distinguished as possible causes of disease by parsing for outlier organisms. Samples with bacterial outliers had significantly depressed alpha-diversity (median 0.58, IQR 0.33-0.62 vs. median 0.94, IQR 0.93-0.95, p<0.001). Potential pathogens were detected in half of samples previously negative by clinical diagnostics, demonstrating increased sensitivity for missed pulmonary pathogens (p<0.001).

**CONCLUSIONS:** An optimized mNGS assay for pulmonary microbes demonstrates significant inoculation of the lower airways of immunocompromised children with diverse bacteria, fungi, and viruses. Potential pathogens can be identified based on absolute and relative abundance. Ongoing investigation is needed to determine the pathogenic significance of outlier microbes in the lungs of immunocompromised children with pulmonary disease.

## BACKGROUND

Last year in the United States, approximately 15,000 children were diagnosed with cancer, 2,000 children underwent solid organ transplantation (SOT), and 2,500 children underwent hematopoietic cell transplantation (HCT) for an increasingly broad set of life-threatening diseases (1–3). While significant progress has been made in improving the safety of anti-neoplastic and transplantation-based therapies, the risk of infectious complications remains high (4). In particular, lower respiratory tract infections in children who have undergone HCT are associated with mortality rates exceeding 45% (5, 6). Current microbiologic diagnostics are inadequate and fail to identify a pathogenic organism in over one third of such cases (7). Autopsy case series have identified previously undetected pulmonary pathogens in 30-50% of pediatric HCT patients, underscoring current diagnostic limitations and the significant mortality associated with undiagnosed pulmonary infections in such patients (8).

Recent advances in metagenomic next generation sequencing (mNGS) have shown promising results in diagnosing neurologic, ocular, and other infections (9–11). However, the identification of fastidious pathogens such as *Aspergillus* and other filamentous molds remain difficult due to thick extracellular matrices and the relatively small inoculum required to induce disease (12–15). Unfortunately, off-the-shelf assays for respiratory biospecimens have proven inadequate to survey the variety of organisms present in thick and mucoid respiratory secretions. As such, unlike the better-characterized microbiomes of the human gastrointestinal tract and nasopharynx, data describing the composition of the pulmonary microbiome are sparse and insufficient to reliably discriminate between health and disease (16).

Therefore, we conducted a pilot study aimed to (1) develop and optimize a highly sensitive mNGS assay capable of detecting pulmonary bacterial, fungal, and viral pathogens clinically relevant to immunocompromised children, and (2) test this mNGS assay on a cohort of immunocompromised children undergoing lower respiratory tract sampling as evaluation for suspected pulmonary infection. We hypothesized that an optimized mNGS assay could improve characterization of the pulmonary microbiome and aid in the identification of potential pulmonary pathogens in this high-risk population.

## METHODS

### Development of Optimized mNGS Assay

Our laboratory has previously characterized high-throughput mNGS protocols for use with human cerebrospinal and intravitreal fluid (9–11). Given the challenges of diagnosing pulmonary infections, we created a set of mock-positive BALs by spiking a pre-determined number of colony forming units (CFUs) of an *Aspergillus niger* clinical isolate into aliquots of BAL with a previously identified *Haemophilus influenzae/Human Adenovirus B* co-infection (**Supplemental Text 1**). *Aspergillus* was chosen as an optimization benchmark given its thick polysaccharide cell wall and extreme clinical importance in this patient population.

### Sample Preparation

We tested nucleic acid extraction conditions by combining 200μL of mock-positive BAL with either 600μL DirectZol, 600μL Lysis Buffer, or 200μL DNA/RNA Shield (Zymo), followed by mechanical homogenization with either 0.1mm or 0.5mm glass bashing beads (Omni) for 2, 5, or 8 cycles of 25 seconds bashing at 30Hz with 60 seconds rest on ice between each cycle (TissuerLyser II, Qiagen). Samples homogenized in DNA/RNA shield also underwent enzymatic mycolysis with either 0.15mg or 0.38mg Proteinase K at 23°C for 30 minutes or 60 minutes (Zymo), or with 0.4mg, 1.2mg, 4mg, or 8mg Yatalase (Takara Bio Inc) at 23°C or 37°C for 60 or 90 minutes. Subsequently, all samples underwent 10 minutes of centrifugation at 4°C and the supernatant was used for paired DNA/RNA extraction (Zymo ZR-Duet DNA/RNA MiniPrep Kit). *Aspergillus* nucleic acid yield was assessed using an orthogonal digital droplet PCR assay with *Aspergillus*-specific primers (ddPCR, **Supplemental Text 2**) (17). Subsequently, sequencing libraries were prepared using the New England Biolabs NEBNext RNA/DNA Ultra-II Library Prep Kit and underwent 125 nucleotide paired-end sequencing on an Illumina HiSeq 4000 instrument (**Supplemental Text 3**).

### Bioinformatics Pipeline

Resultant .fastq files were processed using a previously described pipeline consisting of several open-source components (9, 10). Briefly, reads underwent iterative removal of host (Hg38/PanTro), low quality, low complexity (LZW compression ratio >0.45), and redundant sequences using STAR, Bowtie2, PriceSeqFilter, and CD-HIT-DUP (18–21). The remaining sequences were then aligned to the NCBI non-redundant nucleotide database using GSNAPL for assignment of bacterial, fungal, viral and other taxonomic IDs (22). Microbes were described as potentially pathogenic or typically non-pathogenic based on *a priori* literature review (**Supplemental Text 4**).

### Analysis

To avoid spurious alignments and sequencing artifacts, we only included read pairs where both ends aligned to the same taxonomic ID. The prevalence of each microbe was described ***(1) relative to other microbes in the same sample***, wherein we normalized sequencing reads per million total sequencing reads (rpm); and ***(2) relative to the same microbe in other samples in the cohort***, wherein we normalized ^sequencing reads as the number of standard deviations above or below the mean log_10_-transformed rpm for^ the total cohort (Z-score) (23). Outliers were defined as microbes with RNA alignments that were both abundant within a sample (≥10% of all bacterial reads in a sample, or >1 rpm for fungi and viruses) *and* ≥2 standard deviations above the cohort mean for that particular microbe (Z-score ≥2). A subset of outliers were confirmed via orthogonal assays. The Simpson’s Diversity Index was used to associate the loss of bacterial diversity with the presence of outlier microbes (detailed methods in **Supplemental Text 5**) (24). Sequencing reads are deposited in the NCBI dbGaP (BioProject ID PRJNA437169).

### Patients

To test the optimized mNGS assay, we prospectively collected 41 lower respiratory samples from 34 immunocompromised patients age ≤25 years who underwent clinically-indicated lower respiratory dagnostics between September 2014 and April 2016 at the University of California San Francisco Benioff Children’s Hospital, Indiana University Riley Hospital for Children, and the University of Minnesota Masonic Children’s Hospital (**Supplemental Text 6**). Each sample was labeled as having (1) a pathogen causing respiratory disease, (2) a pathogen not causing respiratory disease, or (3) no pathogen according to interpretation of clinical microbiologic testing as documented by the treating physician. This study was approved by each site’s Institutional Review Board with consent provided by all patients or their surrogates.

## RESULTS

### Development of Optimized mNGS Assay

Iterative optimizations demonstrated that mechanical homogenization of BAL using 0.5mm inert bashing beads for 5 cycles in a DNA/RNA shield without mycolytic enzymes was superior to all other approaches for isolation of microbial nucleic acid (**Supplemental Text 7, Supplemental Figures 1a-d**). At a depth of 25 million reads per 200uL, this reduced the lower limit of detection (LLOD) of RNA sequencing from 59.60 *Aspergillus* CFU (95% CI 37.70-95.36) to 0.42 *Aspergillus* CFU (95% CI 0.12-1.40, paired T-test p<0.001, **Supplemental Figure 2**), demonstrating an approximately 100-fold increased sensitivity for *Aspergillus*. In comparison, the optimized DNA sequencing assay was able to detect as few as 6.13 *Aspergillus* CFU (95% CI 4.16-9.04), suggesting RNAseq to be approximately 10-fold more sensitive than DNAseq for detecting *Aspergillus* nucleic acid in respiratory specimens (paired T-test p<0.001, **Figure 1**). The optimization did not change the detection of *H.influenzae* or *Human Adenovirus B* (T-test p=0.343 and p=0.420, respectively). These results suggest that only the optimized RNAseq protocol had a sufficiently dynamic range to detect *Aspergillus* nucleic acid in quantities analogous to rare growth in culture.

**Figure 1:**
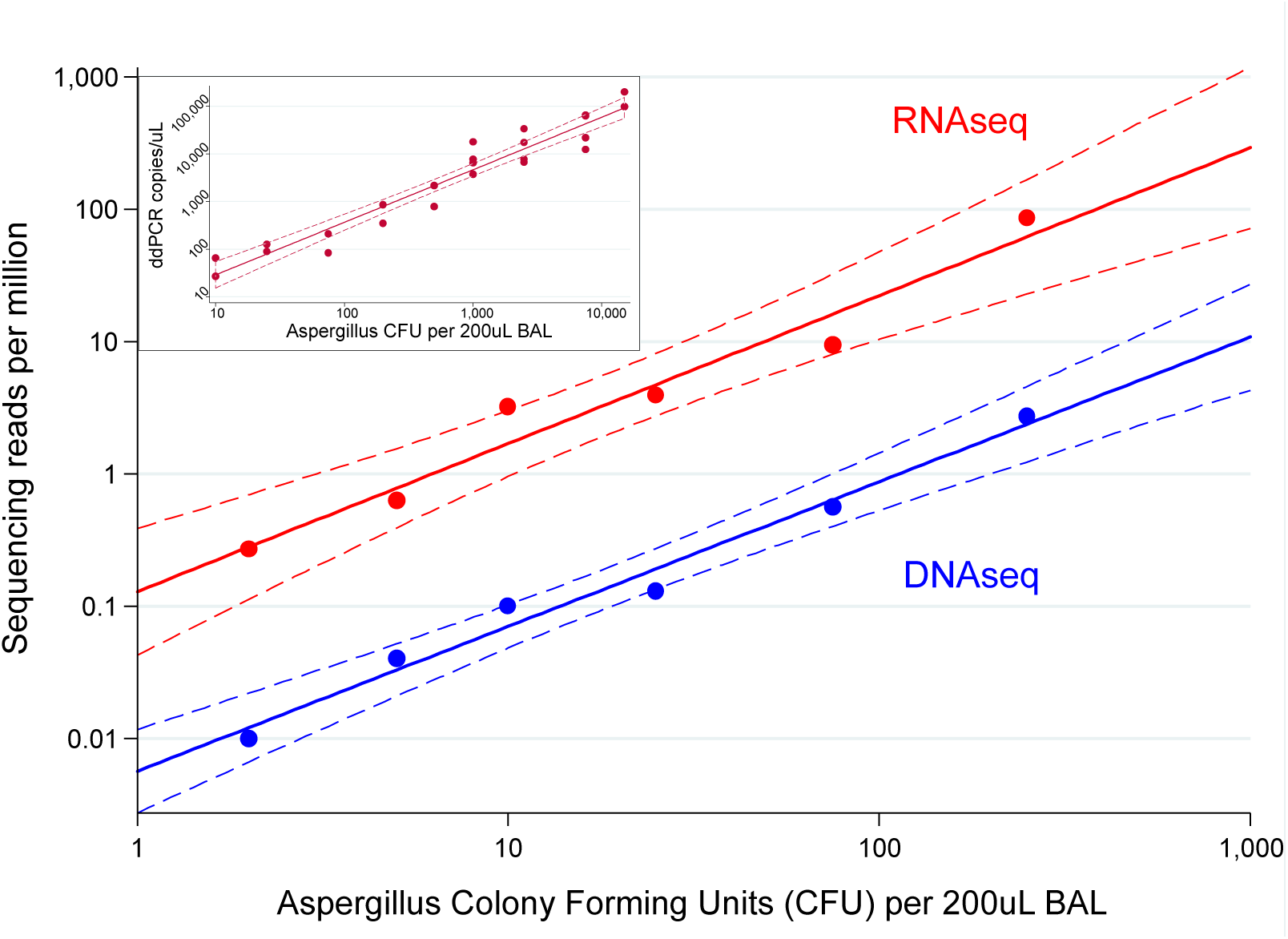
*Aspergillus* Lower Limit of Detection by Optimized Next Generation Sequencing. **Legend: Main Figure:** At the LLOD of RNAseq, the optimized assay was able to detect as few as 0.42 *Aspergillus* CFU (95% CI 0.12-1.40), whereas at the LLOD of DNAseq, the optimized assay was able to detect as few as 6.13 *Aspergillus* CFU (95% CI 4.16-9.04, paired T-test p<0.001). Red data represent RNAseq and blue data represents DNAseq. Dotted lines represent the 95% confidence intervals for each linear regression. **Upper Left Insert:** Parallel detection of *Aspergillus* RNA using ddPCR.

### Application of mNGS Assay

We enrolled 34 patients and collected 41 lower respiratory samples for this pilot project (**Table 1**). Using the optimized extraction protocol described above, we conducted mNGS resulting in 21.2 ±4.2 million reads per sample. After quality filtering and removal of human reads, the remaining reads aligning to bacteria, fungi, and viruses were analyzed further as discussed below. Detailed reports for each patient exist in **Appendix 1**.

**TABLE 1).**
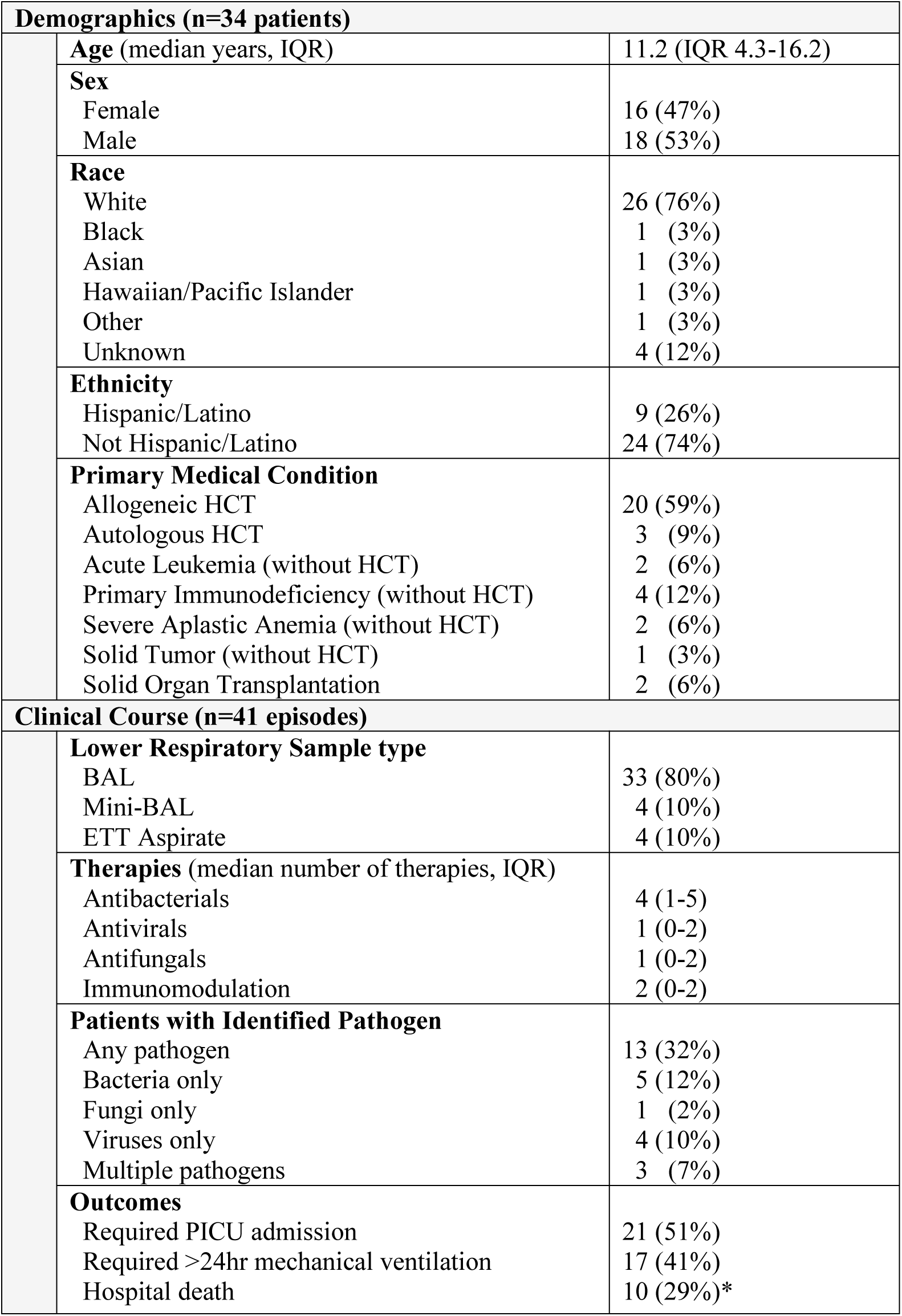
Characteristics of Enrolled Patients. **Legend:** Age at 1^st^ specimen collection. Indications for allogeneic HCT were acute leukemia (12/20), primary immunodeficiency (3/17), severe aplastic anemia (2/17), myeloproliferative/myelodysplastic disorder (2/17), and osteopetrosis (1/17). *Hospital death n=10/34 (29%).

### Bacteria

The vast majority of taxa derived from bacterial alignments were present at low abundance and in quantities similar across all samples in the cohort. Specifically, 95.9% of bacterial genera composed <10% of the total bacterial alignments in each sample and were present at levels within 2 standard deviations of the cohort mean (Z-score <2; **Figure 2a**). These included potentially pathogenic bacteria such as *Escherichia*, *Klebsiella*, *Pseudomonas*, *Staphylococcus*, *Stenotrophomonas*, and *Streptococcus*, which were identified in nearly all patient samples (38, 39, 40, 38, 41, and 40 of 41 samples, respectively).

**Figure 2:**
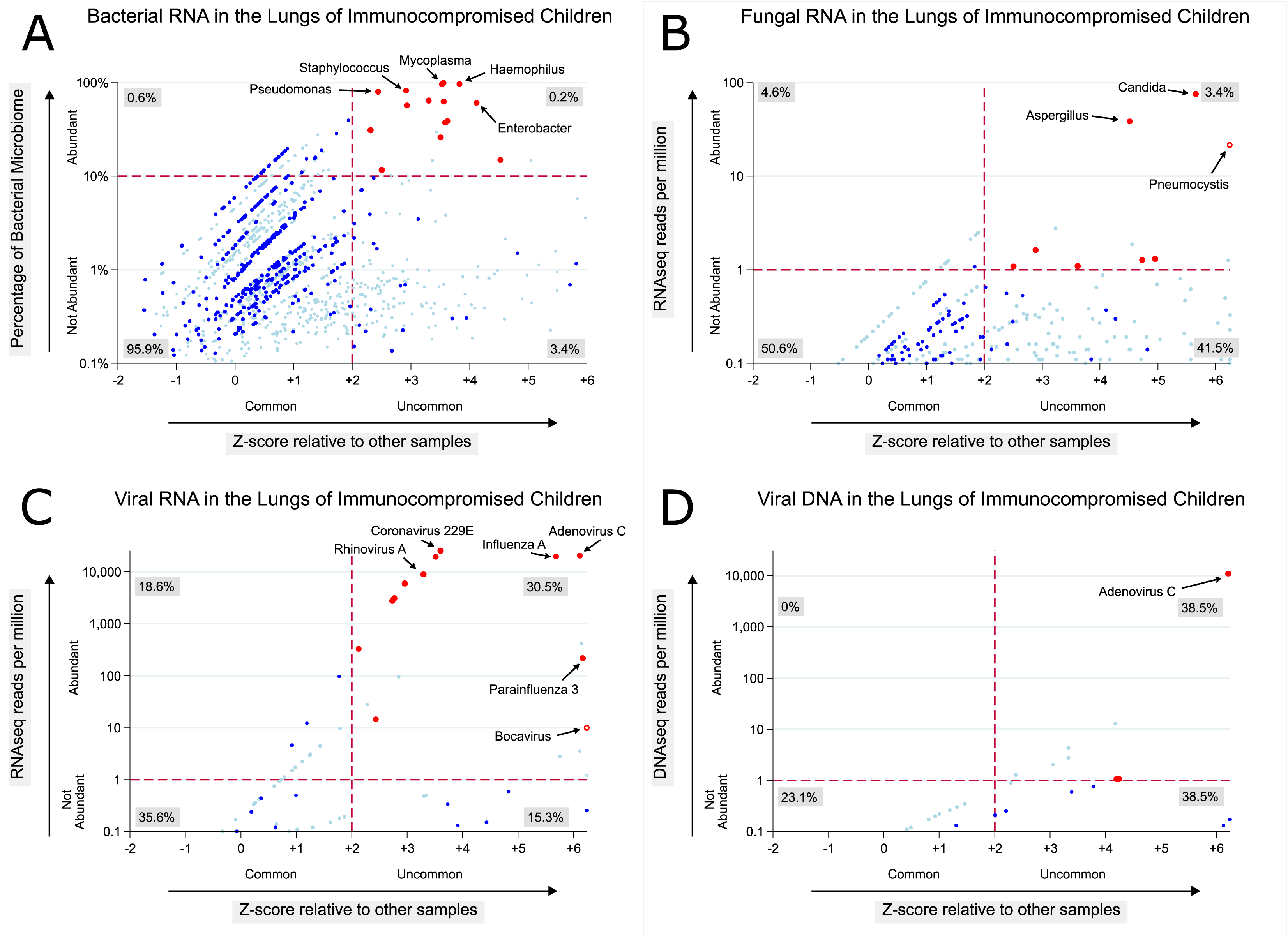
Microbial Alignments Detected in the Lungs of Immunocompromised Children. **Legend:** Red dots represent potentially pathogenic microbes that are both abundant (≥10% of all bacterial reads for bacteria, or ≥1rpm for fungi or viruses) and identified at levels greater than most other samples in the cohort (Z-score ≥2). Hollow red dots indicating *Bocavirus* and *Pneumocystis* are used to indicate organisms observed only once in this cohort. Blue dots represent all other potentially pathogenic microbes; light blue dots represent typically non-pathogenic microbes. Subplots show all bacteria **(A)**, fungi **(B)**, RNA viruses **(C)**, and DNA viruses **(D)** identified across all samples in the cohort. For the purpose of the Z-score calculation, the value of log_10_-transformed reads for undetected microbes was assumed to equal -2, just below the lower limit of detection for our sequencing depth (log_10_[0.01rpm]=-2).

Conversely, only 0.2% of detected bacterial genera composed ≥10% of their respective pulmonary microbiome *and* were present at levels greater than 2 standard deviations above the cohort mean (Z-score ≥2). These outliers included 9 potentially pathogenic bacteria identified in 13/41 patient samples (*Corynebacterium, Enterobacter, Escherichia, Fusobacterium, Haemophilus, Mycoplasma, Pseudomonas, Rothia*, and *Staphylococcus*). These results suggest that parsing pulmonary bacterial reads according to both absolute and relative abundance is an effective way to identify potentially pathogenic bacteria from microbially-rich respiratory samples. Samples with outlier bacterial pathogens had depressed diversity of the bacterial microbiome (median 0.58, IQR 0.33-0.62 vs. median 0.94, IQR 0.93-0.95, p<0.001, **Supplemental Figure 3**), and Simpson’s Diversity Index cutoffs of ≥0.8 or ≥0.9 showed 87.1 % (95% CI 74.7-93.9) and 96.2% (95% CI 79.0-99.4) negative predictive value for the presence of an outlier bacterial pathogen, suggesting that the identification of bacterial dysbiosis may be a useful screen for recognizing possible bacterial infections.

### Fungi

Relative to bacterial alignments, fungal alignments were significantly less prevalent in the cohort. 92.1% of fungal genera were detected at less than a single read per million reads mapped (**Figure 2b**). Approximately half of all identified fungal genera were of low abundance and were present at levels similar to other samples in the cohort (50.6%); an additional 41.5% were of low abundance but were rare in the cohort. Evidence supporting these alignments is limited by the rarity of the reads themselves. Only 3.4% of fungal genera were both abundant and had a Z-score ≥2. These outliers included 7 potentially pathogenic fungi identified a total of 8 times in 7/41 patient samples (*Alternaria, Aspergillus*, *Candida, Cladsoporium, Cryptococcus, Fusarium*, and *Pneumocystis*).

Next, to contextualize the burden of *Aspergillus* in each sample, we estimated the number of *Aspergillus* CFU according to the linear regression associating *Aspergillus* CFU with RNA transcripts derived from our spike-in experiments. *Aspergillus* RNA levels correlated with ≥1 CFU in 14/41 samples, suggesting that the lower airways of immunocompromised children are frequently exposed to quantities of *Aspergillus* RNA consistent with low numbers of CFUs. Of these 14 samples, one patient sample was positive for IPA by both culture and galactomannan assay (**Figure 2b, labeled**). Although the remaining 13 were all culture-negative, 12/13 had received fungicidal or fungistatic anti-*Aspergillus* pharmacotherapy within 48 hours of sample collection, suggesting that empiric antifungal pharmacotherapy significantly confounds the association between *Aspergillus* RNA and growth in culture. While *Aspergillus* RNA sequencing reads did not correlate with growth in culture (T-test p=0.148), they did correlate with BAL galactomannan (regression correlation coefficient 0.74, 95% CI 0.43-1.06, p<0.001 **Supplemental Figure 4**). Samples with *Aspergillus* RNA equivalent to ≥1 CFU had a positive likelihood ratio of 6.0 for having galactomannan ≥0.5 (95% CI 2.5-14.7, p=0.001). These data suggest that while *Aspergillus* growth in culture lacks sensitivity for detecting IPA, mNGS may augment the diagnostic utility of the galactomannan assay.

### Viruses

In this cohort, detected respiratory viruses were dominated by RNA viruses, with *Adenovirus* being the only significant outlying DNA virus. In contrast with bacteria and fungi, a significantly larger portion of RNA alignments to viral genera were present at high abundance, with the majority (30.5%) present in quantities significantly greater than the cohort mean (Z≥2; **Figure 2c, 2d**). Communicable respiratory viruses with abundant alignments were identified in approximately one third of all patient samples (13/41) and included *Adenoviruses A* and *C, Bocavirus, Coronaviruses 229E* and *OC43, Influenzaviruses A* and *C, Parainfluenzavirus 3*, and *Rhinoviruses A* and *C*. Although 4 *Rhinoviruses* were classified as outliers (Z-score ≥2), an additional 3 *Rhinoviruses* were abundant but had Z-scores between +0.92 and +1.78. Additionally, two patients had viral co-infections (*Parainfluenza-3* and *Influenza*-*C*; *Adenovirus-C* and *Rhinovirus-A*).

Approximately half of viral genera were detected below 1 read per million sequencing reads (50.9%). These included additional communicable respiratory viruses at low abundance in 15% of the cohort (6/41). Although there were no cases of clinically suspected herpesvirus pneumonitis, *Epstein-Barr virus*, *Cytomegalovirus*, *HHV-6*, and *HHV-7* were identified in low abundance in 9/41 samples. Viral genera of uncertain or unlikely pathogenicity were also identified in 21/41 samples and included *Papillomaviruses*, *WU* and *KI Polyomaviruses*, and *Torquetenoviruses* (2, 5, and 18 of 41 samples, respectively).

### Comparison to Clinical Testing

Clinical testing identified causative pathogens in 36.6% of samples (n=15), and each of these were concordantly identified by mNGS, although 4 were not classified as outliers (**Figure 3**). mNGS also identified 4 previously undetected potential co-pathogens in this group (*Bocavirus, Influenza-C*, and *Coronavirus* twice). Clinical testing identified non-causative pathogens in 12.2% of samples (n=5); 4 were concordantly identified by mNGS and none were ranked as outliers. However, abundant transcripts aligning to *Human Coronavirus 229E* provided an alternative explanation for pulmonary disease in two of these patients. Clinical testing did not identify any pathogens in 51.2% of samples (n=21). Here, mNGS was able to identify statistically outlying potential pathogens in 10/21 cases. These potential pathogens included a variety of bacteria (ie: *P.aeruginosa*, *E.cloacae, M.pneumoniae*), fungi (ie: *C.glabrata*), and viruses (ie: *Rhinovirus-A*, *Influenza-A)*. Among patients whose clinical testing did *not* identify a causative pathogen, we observed a non-significant trend towards greater mortality in patients with an outlier pathogen detected on mNGS (5/11 vs. 1/12, Fisher p=0.068).

**Figure 3:**
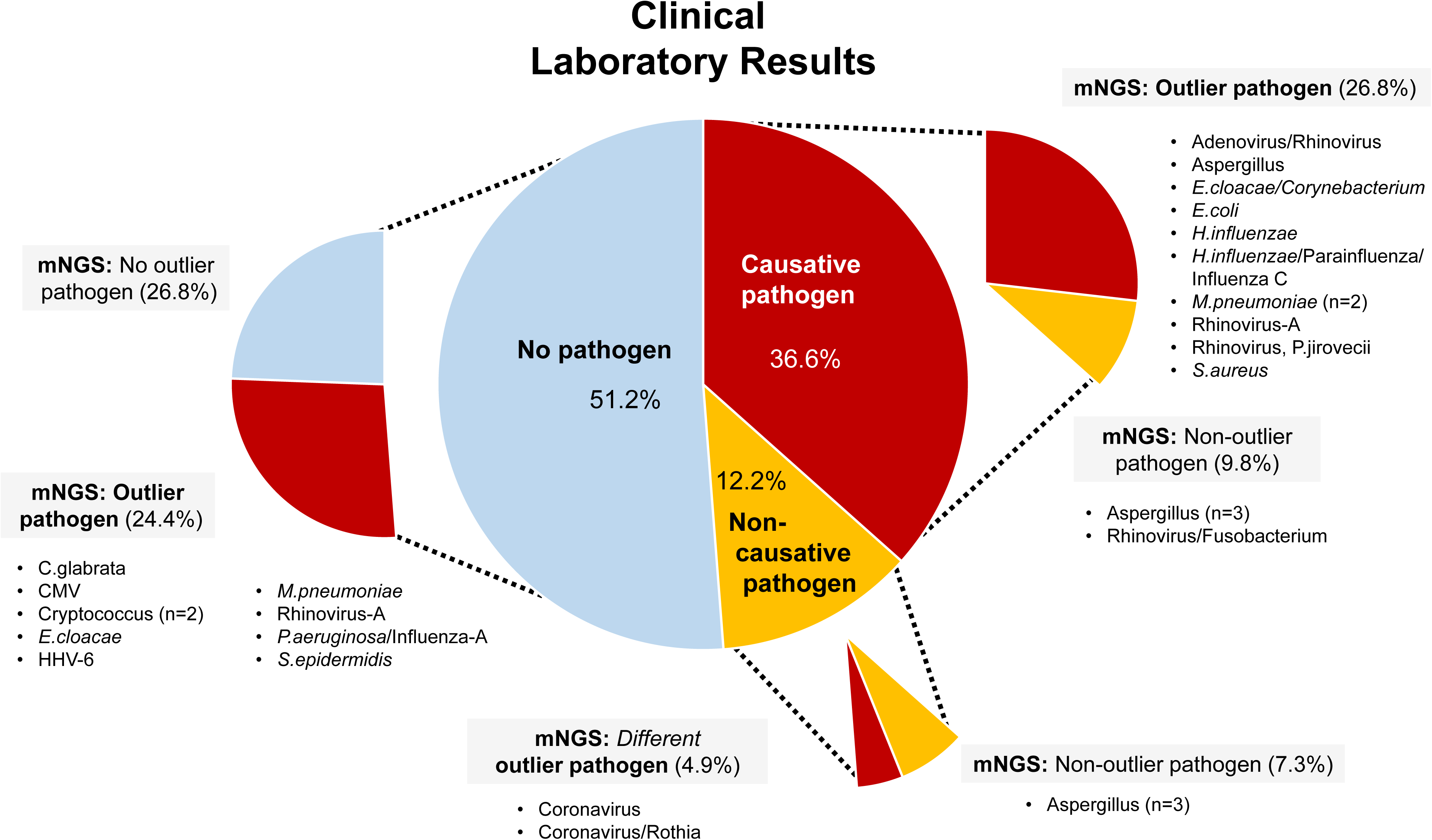
Comparison of Clinical Laboratory Results vs. mNGS Results. **Legend:** Clinical laboratory results were determined by review of medical charts. Pathogens reported by clinical laboratories but determined by the treating clinical team to *not be* causative of the patient’s underlying respiratory disease were labeled as unlikely pathogens.

### Orthogonal Validation

Organisms detected by mNGS but missed during clinical testing were validated with commercially available CLIA-approved assays, including multiplex respiratory viral PCR, *Mycoplasma*-specific PCR, and 16S bacterial and 28/ITS fungal amplicon sequencing (**Supplemental Text 5**). All validation tests were concordant with mNGS with one exception: 28S/ITS amplicon DNA sequencing failed to identify *C.glabrata* in Sample 37. For this sample, we confirmed the presence of this organism with 3 separate species-specific RT-PCR primer sets followed by Sanger sequencing in-house, confirming the initial mNGS report of *C.glabrata*. As there are numerous complementary bioinformatics approaches for analyzing metagenomic datasets, we tested the reproducibility of our approach by re-analyzing our data using a parallel method developed in-house that weights organisms by both abundance and concordance of paired RNA and DNA sequencing reads without Z-scores (**Supplemental Table 1**). The results of this analytical approach and our primary analytical approach showed strong agreement. In sum, these data demonstrate that mNGS has significantly greater sensitivity for detecting potential pulmonary pathogens than current clinical diagnostics (McNemar’s p<0.001).

## DISCUSSION

In this study, we developed an optimized mNGS assay with adequate sensitivity to identify bacteria, fungi, and both RNA and DNA viruses within the lower respiratory tract of immunocompromised children. In doing so, we identified a rich molecular portrait of the pulmonary microbiome in this vulnerable patient population. Further, by comparing the quantity of microbial nucleic acid to other microbes within a sample and to other samples within the cohort, we were able to identify outlying potential pathogens in approximately half of previously-negative samples.

Due to inherent challenges in sampling the lower respiratory tract, the pulmonary microbiome was not one of the original sites sampled in the 2008 Human Microbiome Project and data regarding pulmonary microbial communities in health and disease have lagged decades behind similar analyses of human intestinal, cutaneous, and nasopharyngeal microbiomes (27). In this study, we detected that many pathogenic bacteria such as *Pseudomonas* and *Streptococcus* are ubiquitous and hence their abundance needs to be contextualized by cohort-specific norms. By using cohort-specific Z-scores, we provide a template for discriminating normal from abnormal microbial burden in the lungs of immunocompromised children. For example, 100% of samples had detectable *Pseudomonas* RNA, but only Sample 29 had detectable *Pseudomonas* RNA more than 2 standard deviations above the cohort mean (**Appendix 1**).

While prior metagenomic sequencing assays for human biospecimens have demonstrated excellent detection of viral and bacterial nucleic acid, they have lacked sensitivity for detecting fastidious organisms such as filamentous mold (12, 13). This study confirms the need for aggressive mechanical homogenization in stabilizing media in order to detect molds such as *Aspergillus* while simultaneously preserving the detection of bacteria and viruses (25). Despite this, transcripts aligning to fungi remained significantly less ubiquitous than transcripts aligning to bacteria, perhaps due to relative organism abundance. Although the majority of commercial sequencing assays use DNA, our data demonstrate that RNA sequencing is >10 times as sensitive as DNA sequencing for the detection of such fastidious organisms, which we speculate may be due to high transcriptional activity of regions such as rRNA genes in active organisms (26).

Surprisingly, *Aspergillus* RNA equivalent to ≥1 CFU was detected in 34.1% of samples and no water controls, suggesting that the lungs of immunocompromised children are frequently exposed to low levels of *Aspergillus.* The 100-fold increase in *Aspergillus* nucleic acid after optimized mechanical homogenization suggests that the origin of the *Aspergillus* RNA is likely from intact fungal organisms in the lower respiratory tract of immunocompromised children, rather than from migration of extracellular nucleic acid down the respiratory tree. However, as only 10% of samples originated from patients with suspected IPA, we speculate that patient-specific factors such as immune reconstitution, alloreactive inflammation, and impaired mucociliary clearance are as important as inoculum exposure in determining which child might develop IPA (28). Although *Aspergillus* growth in culture appeared to be confounded by the frequent use of prophylactic and empiric antifungals, the quantity of *Aspergillus* sequencing reads was correlated with BAL galactomannan (29, 30). As nutrient availability and the kinetics of *Aspergillus* growth may affect both galactomannan release and bioavailability of *Aspergillus* nucleic acid, we advocate that RNA/DNA-based assays might complement but should not replace antigen-based assays in the diagnosis of IPA (31, 32). While our mNGS assay was highly sensitive for *Aspergillus* detection, specificity for differentiating infection from colonization was limited; future studies pairing mNGS with host susceptibility and immune function may improve discrimination of colonization and invasive mycosis (33–36).

Consistent with emerging data in pediatrics, this work also detected herpesviruses such as *CMV* and *HHV-6* in low abundance across many children in the cohort (37, 38). Interestingly, a number of typically non-pathogenic viruses such as *Torquetenovirus* and *KI Polyomavirus* were present in over half the cohort. While data supporting the direct pathogenicity of these viruses are lacking, they may be a marker of immune function and a predictor of post-transplant infectious and alloimmune complications, including lung injury in general (39–41). Future studies aimed at characterizing longitudinal changes in the pulmonary virome of immunocompromised children are warranted.

By characterizing the distribution of pulmonary microbes within the cohort, we were able to identify statistically outlying potential pathogens in half of previously negative samples. Most cultivable bacteria and fungi that were not detected clinically were isolated from patients who had already received antimicrobial therapies, highlighting the importance of culture-independent techniques (16). In addition, several pathogenic viruses were not detected clinically due to omission from multiplex PCR assays (*Human coronavirus, Human bocavirus, Influenzavirus-C*) (42, 43). Unfortunately, a number of pathogens detected by mNGS currently lack targeted therapeutics, and as such, the development of pharmacologic and cell-based therapies remains a priority in this field (44–46). In addition, while this study suggests that many idiopathic pulmonary complications may indeed be triggered by infections, half of samples without pathogens identified by clinical testing were concordantly negative on mNGS, emphasizing the ongoing clinical significance of non-infectious pulmonary complications in this population. Future clinical validation of mNGS may demonstrate utility in safely excluding infections and allowing amplified immunosuppression in patients with suspected non-infectious pulmonary syndromes such as hepatic veno-occlusive disease, thrombotic microangiopathy, and pulmonary graft-versus-host disease (47, 48).

### Strengths and limitations

Our study has several strengths. First, we optimized the identification of *Aspergillus* RNA and contextualized *Aspergillus* sequencing reads by comparing with both growth in culture and BAL galactomannan. Second, we proposed a logical analytical framework that ranks organism abundance both within a sample and relative to other samples; this framework allowed us to rapidly exclude more than 95% of identified bacteria. Third, we provide, to our knowledge, the first evaluation of the pulmonary microbiome in immunocompromised children.

This pilot study has several limitations. First, while using standard Z-scores was useful in de-emphasizing commonly abundant organisms (ie: *S.pneumoniae*), it may have overvalued uncommon pathogens with less abundant transcripts (ie: *CMV*) or undervalued frequently encountered pathogens (ie: *Rhinovirus*). Additional larger studies aimed at quantifying pulmonary microbial communities in immunocompromised children will naturally improve the value of Z-score analysis. Second, as with all mNGS assays, the identification of microbial nucleic acid does not directly confirm the presence of viable, live organisms, does not directly implicate that microbe as a contributor to pulmonary disease, and does not exclude less abundant organisms as potential contributors to pulmonary disease. In order to optimize patient outcomes, we advocate for ongoing multidisciplinary collaboration between intensivists, immunologists, oncologists, transplant physicians, infectious disease physicians, pulmonologists, bioinformaticians, and basic scientists.

## CONCLUSIONS

In summary, we present a mNGS assay optimized to evaluate the pulmonary microbiome and detect potential pathogens in immunocompromised children. This assay is highly sensitive, reveals a rich bacterial, fungal, and viral pulmonary microbiome, and further identified potential pathogens in half of previously negative samples. As such, advanced organism detection offers the potential for early implementation of targeted therapy and the possibility for improved clinical outcomes in immunocompromised children. We invite the scientific and clinical community to participate in an ongoing multicenter collaborative clinical trial aimed at further refining this emerging technology (https://clinicaltrials.gov/ct2/show/NCT02926612).

## DESCRIPTOR

10.11 Pediatrics: Respiratory Infections

## SUPPLEMENTAL DATA

This article has an online data supplement. Non-human reads from each patient are deposited in the NCBI SRA (BioProject ID PRJNA437169).

AUTHOR CONTRIBUTIONS
*Study concept and design*: MSZ, CCD, AS, JDL
*Acquisition of data:* MSZ, CCD, MYM, KI, NPL, MEM, GDC, LEF, CMW, JRH, MES, SM, KS, AM, GAY, AS, JDL
*Analysis and interpretation of data*: MSZ, CCD, MYM, EDC, CL, KK, EDC, SM, JDL
*Drafting of the manuscript*: MSZ, CCD, MYM, JDL
*Critical revision of the manuscript for important intellectual content*: MSZ, CCD, MYM, KI, NPL, MEM, GDC, LEF, CMW, JRH, MES, EDC, CL, KK, EDC, SM, KS, AM, GAY, AS, JDL
*Statistical analysis*: MSZ, MYM, KK, JDL
*Administrative, technical, or material support:* MSZ, CCD, SM, AS, JDL
*Study supervision*: MSZ, CCD, AS, JDL
*Approval of final manuscript:* MSZ, CCD, MYM, KI, NPL, MEM, GDC, LEF, CMW, JRH, MES, EDC, CL, KK, EDC, SM, KS, AM, GAY, AS, JDL

